# Nuclear genetic background influences the phenotype of the *Drosophila tko^25t^* mitochondrial protein-synthesis mutant

**DOI:** 10.1101/2023.02.06.527322

**Authors:** Howard T. Jacobs, Tea Tuomela, Päivi Lillsunde

**Author notes:** Corresponding author: Howard T. Jacobs, Faculty of Medicine and Health Technology, FI-33014 Tampere University, Finland.

## Abstract

The *Drosophila tko^25t^* point mutation in the gene encoding mitoribosomal protein S12 produces a complex phenotype of multiple respiratory chain deficiency, developmental delay, bang-sensitivity, impaired hearing, sugar and antibiotic sensitivity and impaired male courtship. Its phenotypic severity was previously shown to be alleviated by inbreeding, and to vary with mitochondrial genetic background. Here we show similarly profound effects conferred by nuclear genetic background. We backcrossed *tko^25t^* into each of two standard nuclear backgrounds, Oregon R and *w^1118^*, the latter used as recipient line in many transgenic applications requiring selection for the *white* minigene marker. In the *w^1118^* background, *tko^25t^* flies showed a moderate developmental delay and modest bang-sensitivity. In the Oregon R background, males showed longer developmental delay and more severe bang-sensitivity, and we were initially unable to produce homozygous *tko^25t^* females in sufficient numbers to conduct a meaningful analysis. When maintained as a balanced stock over 2 years, *tko^25t^*flies in the Oregon R background showed clear phenotypic improvement though were still more severely affected than in the *w^1118^* background. Phenotypic severity did not correlate with the expression level of the *tko* gene. Analysis of tko25t hybrids between the two backgrounds indicated that phenotypic severity was conferred by autosomal, X-chromosomal and parent-of-origin dependent determinants. Although some of these effects may be *tko^25t^*-specific, we recommend that, in order to minimize genetic drift and confounding background effects, the genetic background of non-lethal mutants should be controlled by regular backcrossing, even if stocks are usually maintained over a balancer chromosome.

## INTRODUCTION

The *Drosophila* bang-sensitive mutant *tko^25t^*, harboring a point mutation in the gene encoding mitoribosomal protein S12 (Royden et al. 1987; Shah et al. 1997), was originally classified as semilethal (Judd et al. 1972). It was subsequently maintained by ourselves and others, both as an inbred stock and over an FM7 balancer chromosome. The major features of the organismal phenotype, in addition to bang sensitivity, are delayed larval development, deafness (non-responsiveness to sound), impaired male courtship and hypersensitivity to antibiotics targeting the mitochondrial translation system (Toivonen et al. 2001). In an earlier study we reported that the inbred, laboratory-maintained *tko^25t^* stock manifested a milder phenotype than originally described, but that a more severe phenotype could be restored by backcrossing the mutation into two different standard nuclear backgrounds (Canton S and Oregon R) to create a hybrid (Jacobs et al. 2004). Subsequently we have explored various genetic and environmental manipulations that further modified the *tko^25t^* phenotype. Individual lines established from the original *tko^25t^* Canton S/Oregon R hybrid, and selected at each generation for fastest development, showed varying degrees of suppression of the mutant phenotype. The three most pronounced suppressors were found to harbor segmental duplications of the region of the X chromosome encompassing the mutated *tko* gene (Kemppainen et al. 2009).

Mitochondrial genetic background was also shown to be a key determinant of *tko^25t^* phenotype: in some mtDNA backgrounds, notably those previously infected with the endosymbiont *Wolbachia*, partial suppression was found (Chen et al. 2012), whilst another mtDNA haplotype isolated from the wild was synthetically lethal with *tko^25t^* (Salminen et al. 2019). Here we attempted a similar analysis, but of the effect of nuclear background, keeping mtDNA haplotype and the *tko^25t^* mutation itself unchanged. Without imposing any kind of deliberate selection, we obtained divergent outcomes, pointing to the genetic complexity of nuclear-mitochondrial interactions in a metazoan.

## MATERIALS AND METHODS

### *Drosophila* culture and crosses

Flies were cultured in standard high-sugar medium (Fernandez-Ayala et al. 2009): 1 % agar, 3% glucose, 1.5% sucrose, 3% treacle (Tate & Lyle, UK), 3.5% dried yeast, 1% soy flour, 1.5% maize flour, plus 0.1% Nipagin M (Sigma) and 0.5% (v/v) propionic acid (JT Baker) as antimicrobials, on a 12 h/12 h light-dark cycle at the temperatures indicated in figures and legends (25 °C where not otherwise stated). For long-term maintenance fly stocks were kept at 18 °C. The *tko^25t^* mutation was maintained over an FM7 balancer chromosome. *tko^25t^* flies were generated for experimental analysis by crossing *tko^25t^* / FM7 females with *tko^25t^* / Y males, with an initial cross of *tko^25t^*/ FM7 females with FM7 / Y males to generate the latter, where needed.

### Eclosion timing and bang-sensitivity

Eclosion timing was measured, as previously (Kemppainen et al. 2016), using replicate sets of vials (n ≥ 3). Note than mean eclosion times shown in figures for each sex and genotype are the mean of the replicate vials in a given experiment. Numbers of adult progeny are indicated in figures and/or legends. Bang-sensitivity was measured as previously (Toivonen et al. 2001): individual flies were transferred under CO_2_ anesthesia from a food vial into a single empty vial with a bung. After collection, flies were left for a minimum of 1 hour to recover from CO_2_ exposure. Each fly was subjected to 10 seconds of vortexing on a laboratory vortex-mixer at maximum speed. The vial was then tapped so that the paralyzed fly fell to the bottom of the vial on its back. Each fly was continuously observed for up to 300 s until it recovered a normal standing posture, with the elapsed time scored as the recovery time. Recovery time was arbitrarily scored as 300 s for any fly that did not resume a normal standing posture during the experiment, and as 0 s for any fly that did not visibly become paralyzed at all.

### Single-fly genomic DNA extraction

Individual flies in a 0.5 ml tube were mashed for 5-10 s with a plastic pipette tip in 50 μl of SB (10 mM Tris-HCl, 1 mM EDTA, 25 mM NaCl, 200 μg/ml Proteinase K, pH 8.2). After incubation at room temperature for 20-30 min the Proteinase K was inactivated by heating to 95 °C for 1-2 min and the mix used immediately for PCR.

### Genotyping

The presence of the *tko^25t^*mutation was verified by PCR amplification of a 1.49 kb fragment of the *tko* gene from single-fly genomic DNA, using primer Tko-51 (5′-GAACAAAAAACTACTGAACAAAACTCC-3′) and Tko-31 (5′-CATTTGAACAACGTGATTAGGAAGT-3′), followed by Sanger sequencing of the agarose gel-purified product using primers tko_TT_L1 (5′-GTGCTTTATTGATTTCGAGCGATCT-3′) and tko_TT_R1 (5′-ACTATTGGCTCTTCTTGACGACGTG-3′).

### Expression analysis by qRTPCR

Total RNA was extracted and *tko* RNA levels relative to *RpL32* were measured by quantitative RTPCR as previously using the SYBR Green method (Bahhir et al. 2019) using the following oligonucleotide primers (all shown 5′ to 3′): for *tko*, tko_L1_qPCR – CGACGGCTGGTACTACAAAC and tko_R1_qPCR – GGTCTCATAGCTGCACTGGA and for *RpL32*: RpL32_F – AGGCCCAAGATCGTGAAGAA and Rpl32_R – TGTGCACCAGGAACTTCTTGAA.

### Courtship analysis

Courtship was analyzed essentially as previously (Toivonen et al. 2001), in batches of 6 pairs of adult male and virgin female 3 day-old flies of a given genotype. For experiments at different temperatures, virgin flies were kept at the experiment temperature until the experiment was started. Flies were transferred into individual mating chambers by drawing them from culture vials one by one using a depressurized plastic bottle and releasing into the chamber. With the female and male initially separated, mating chambers were placed in an incubator at the specified temperature. Flies were allowed to recover from the transfer for one hour before allowed to interact with the opposite sex and then inspected under a stereo microscope at 15-minute intervals during each hour-long experiment to determine how many pairs copulated. Mean copulation frequency on a per batch basis was derived from n > 5 batches of each genotype studied

### Statistical analysis

Data were analyzed, where indicated, either by Student’s *t* test or by one-way ANOVA using a standard online tool (https://astatsa.com/OneWay_Anova_with_TukeyHSD/). Recovery times in the bang-sensitivity assay were not normally distributed: therefore, raw data only are presented in the figures.

## RESULTS AND DISCUSSION

### Backcrossing of *tko^25t^* into standard backgrounds

The *tko^25t^* mutation was backcrossed over ≥10 generations according to the scheme outlined in Figure 1 into each of two standard nuclear backgrounds, Oregon R, used in laboratories worldwide since 1925 – see (Lindsley and Grell, 1968), and *w^1118^*(Hazelrigg et al. 1984), widely used for selection of transgenic lines using the *mini-white* marker, which confers colored eyes in a white-eyed background. After back-crossing, several lines in each background were maintained over a standard X-chromosomal balancer, FM7 (Merriam, 1968). For each background, we established multiple independent back-crossed lines.

**Figure 1:**
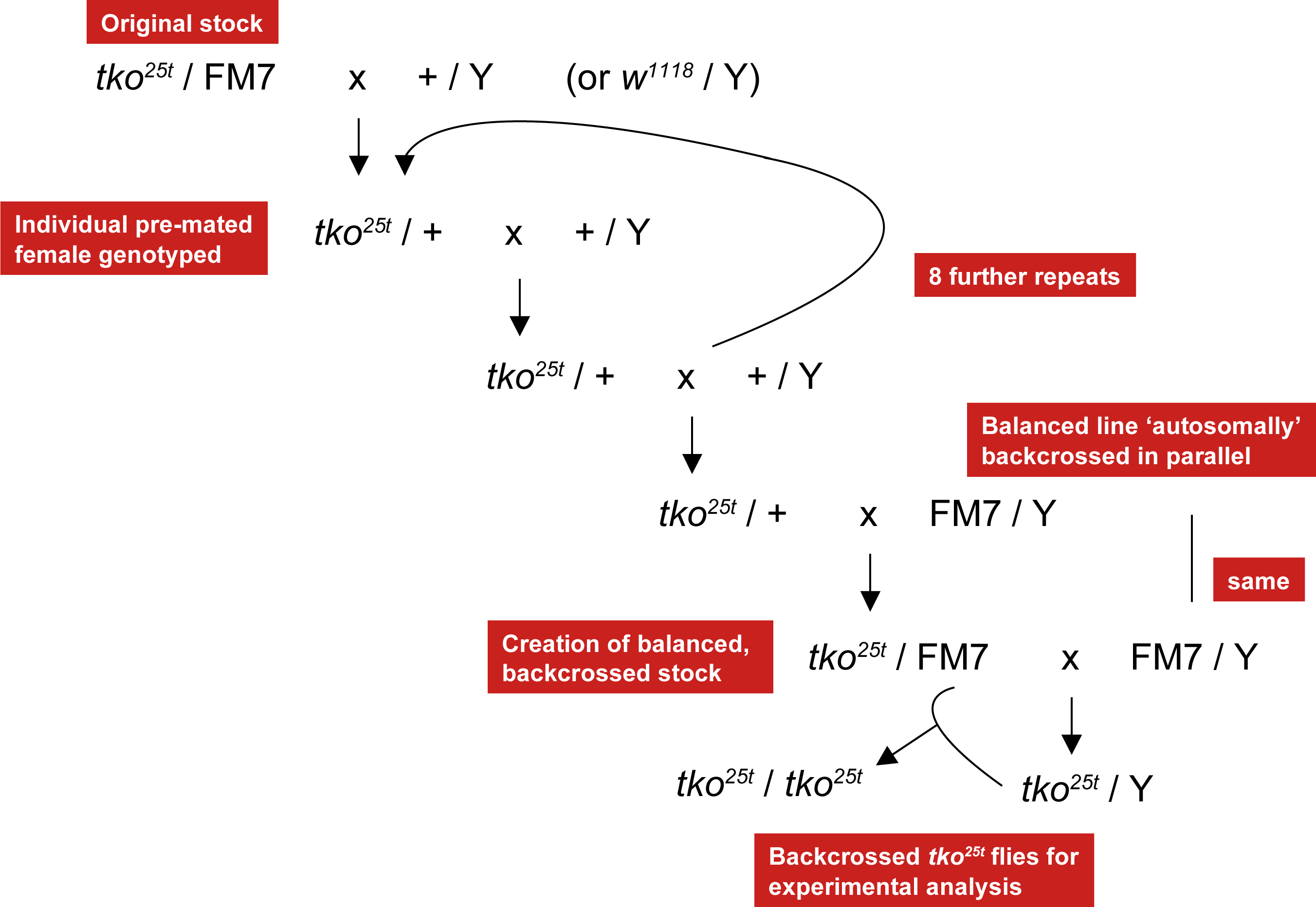
Backcrossing scheme for *tko^25t^*. At each backcross generation, heterozygous (*tko^25t^ /* +) females were identified by single-organism genomic DNA extraction, PCR and sequencing, after pre-mating. Progeny from wild-type females was discarded. Several parallel lines were established in each backcross, both to the *w^1118^* and to the Oregon R backgrounds. The FM7-balanced, ‘autosomally backcrossed’ lines were established in parallel.

### Outbred *tko^25t^* lines manifest background-dependent phenotypes

All of the outbred *tko^25t^* lines showed a more severe phenotype than the lab-maintained *tko^25t^* stock, in terms of both developmental delay and bang-sensitivity. The *tko^25t^* phenotypes in the two backgrounds were also phenotypically distinct (Fig. 2), although three independent back-crossed lines in the Oregon R background were similar to each other, as were three such lines in the *w^1118^* background (Fig. 2). In initial crosses to generate *tko^25t^* males from the balanced outbred stocks (see legend to Fig. 2), we observed that very few mutant males eclosed in the Oregon R background at the standard culture temperature of 25 °C, resembling the semilethality first reported for the mutation (Fig. 2A), whereas the output of mutant males in the *w^1118^* background was very close to the expected 25% (Fig. 2A). The few *tko^25t^* males in the Oregon R background that eclosed were very delayed and extremely bang-sensitive, although there were too few from each line to generate meaningful numbers. For the *w^1118^* background, developmental delay at 25 °C was around 5 days, whereas our original *tko^25t^* stock showed a developmental delay of only 2-3 days (Fig. 2B). We found that more substantial numbers of *tko^25t^* males in the Oregon R background eclosed at room temperature 21-22 °C, although they were still very bang-sensitive and did not mate, making it impossible to generate *tko^25t^*females in this background. We were, however, able to generate sufficient numbers of mutant males to compare developmental delay (Fig. 2C) and bang-sensitivity (Fig. 2D) in the two backgrounds.

**Figure 2:**
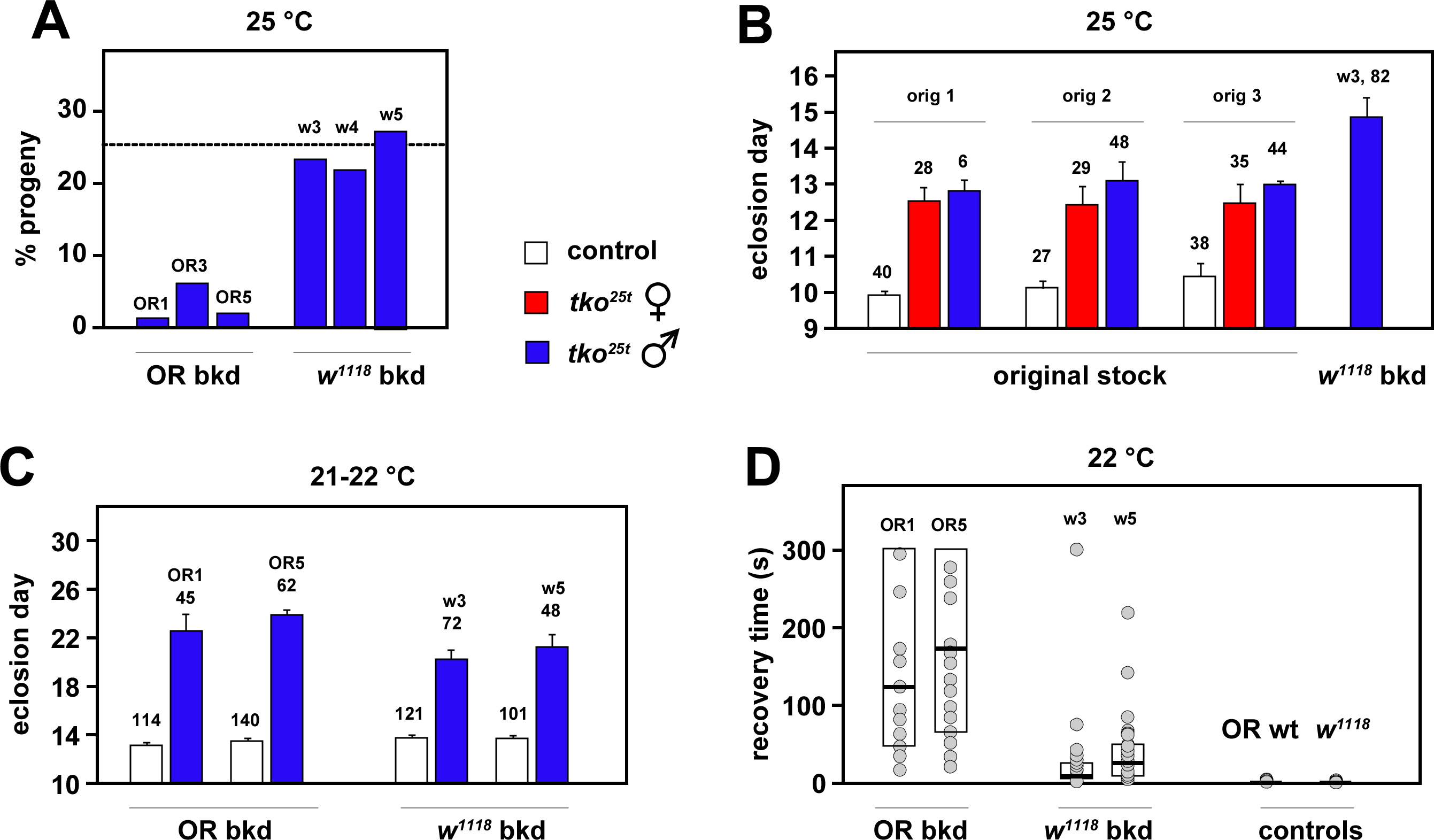
**Phenotype of *tko^25t^* in different nuclear backgrounds**. (A) Proportion of *tko^25t^* males amongst eclosing progeny from crosses of the type *tko^25t^* / FM7 x FM7 / Y, conducted at 25 °C, after back-crossing of *tko^25t^* into the Oregon R background (OR bkd), to create lines OR1, OR3 and OR5 as indicated, or into the *w^1118^* background, to create lines w3, w4 and w5. The dotted line indicates the expected proportion of *tko^25t^* / Y progeny (25%). n = 208 (OR1), 295 (OR3), 314 (OR5), 363 (w3), 330 (w4) and 338 (w5) total eclosed adults, from 4 individual vials in each case. In the original, lab-maintained *tko^25t^*stock, the percentage of mutant males in similar crosses without other balancers was always close to the expected 25%, i.e. similar to the findings for *tko^25t^* backcrossed into *w^1118^*. The output of males in each of the two background control strains in equivalent crosses was routinely the expected ∼50%. (B) Eclosion times (means ±SD) for *tko^25t^*flies generated in multiple crosses using the original *tko^25t^*stock. In the three experiments shown (orig 1, orig 2, orig 3), mothers were *tko^25t^* / FM7 and carrying different autosomal balancers (respectively, CyO, TM3-Sb and TM3-Ser) with the scored *tko^25t^*progeny free of all balancer markers, some of which are known to interact with *tko^25t^* (see Fig. S2 of George et al.2019, which presents some of the same data). The number of adult flies of each genotype/sex analyzed is indicated above the applicable column. n = 5 replicate vials in all cases except for line w3, where n = 4 replicate vials. All controls were *tko^25t^* / FM7 heterozygotes in the same background which, in both back-cross backgrounds, gave indistinguishable eclosion times as wild-type males in the same background (Fig. S1A). (C) Eclosion times (means ± SD) for similar crosses as in (A), but carried out at room temperature (21-22 °C). The number of adult flies of each genotype/sex analyzed is indicated above the applicable column. n = 5 replicate vials in all cases. (D) Bang-sensitivity assay (recovery times) for progeny flies of the indicated genotypes from the crosses in (C), assayed at 22 °C. n = 35 (OR1), 38 (OR5), 38 (w3) and 29 (w5) *tko^25t^* mutant males, and 119 (OR) and 132 (*w^1118^*) control males. Box plots show 25^th^ and 75^th^ percentiles of each distribution and medians (bold lines). Note that the scatter plots are only indicative, since many data points are coincident or almost completely overlapping on this scale: for full details of source data see Table S1.

### Outbred *tko^25t^* lines maintained as balanced stock show phenotypic alleviation

The *tko^25t^* mutation in the outbred lines was maintained for two years over an FM7 balancer chromosome with no intentional selection. When retested, the phenotype of the lines in both backgrounds was substantially milder than in the initial tests. In the Oregon R background the mutation was no longer semilethal, and mutant males were now able to mate and give offspring. We were able to obtain sufficient numbers of mutant flies of both sexes and both backgrounds to reliably measure developmental time (Fig. 3A) and bang-sensitivity (Fig. 3B) at 25 °C (Fig. 3). Flies outbred to the *w^1118^*background still showed a milder phenotype than those in the Oregon R background, notably for bang-sensitivity (Fig. 3B), but were also less severely affected than when initially tested, immediately after the completion of the backcrosses. Developmental delay of *tko^25t^*flies at 25 °C was 3-4 d in the *w^1118^*background and 4-5 d in the Oregon R background (see examples in Fig. 3), with males always more delayed than females.

**Figure 3:**
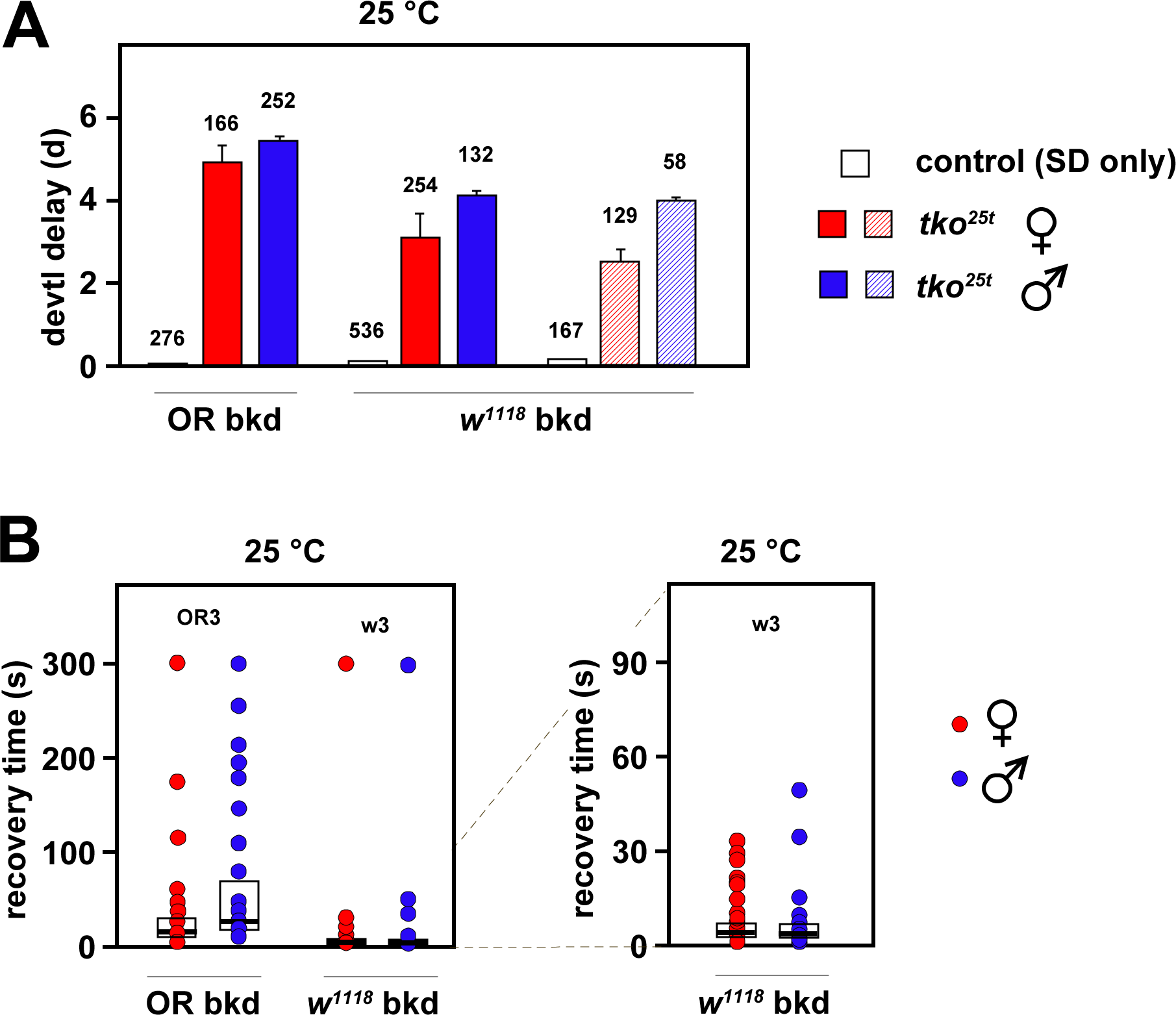
Phenotype after 2 years in culture of *tko^25t^* backcrossed into different nuclear backgrounds. (A) Developmental delay (days, d, means ± SD) and (B) bang-sensitivity recovery times (box-plot nomenclature as in Fig. 2) for *tko^25t^*flies and controls of the indicated back-crossed lines, after 2 years in culture as balanced stocks. Crosses of the type *tko^25t^* / FM7 x *tko^25t^* / Y were conducted at 25 °C, with controls being *tko^25t^* / FM7 heterozygotes. In (A) a repeat experiment is shown for the w3 line (hatched rectangles), due to the large error bars seen for mutant females in the primary experiment. The number of adult flies of each genotype/sex analyzed is indicated above the applicable column. n = 3 replicate bottles in all cases. In (B) the same data for *tko^25t^* flies in the *w^1118^* background are shown at two different scales, as indicated by the dotted lines, with the 300 s data points omitted, for clarity, in the expanded-scale graph (right-hand panel). n = 70 (OR3 females), 46 (OR3 males), 104 (w3 females) and 85 (w3 males). Note that, as for Fig. 2D, the scatter plots are only indicative, since many data points are coincident or almost completely overlapping on both scales: for full details of source data see Table S2. Control flies (see Fig. 2D) were not bang-sensitive.

### *tko^25t^* phenotype and *tko* expression are crudely correlated

In a previous study (Kemppainen et al. 2009) no strict correlation was found between the expression level of the mutant *tko* gene and the *tko^25t^* phenotype, although duplication of the mutant gene in its natural chromosomal milieu did provide a partial phenotypic rescue (Kemppainen et al. 2009). In the present study it was clear that the phenotype of developmental delay and bang-sensitivity was more pronounced in mutant males than mutant females, in the Oregon R background compared with the *w^1118^* background and in mutant flies cultured at 25 °C than at 21-22 °C. To re-examine the influence of *tko* expression level on phenotype we used qRTPCR to analyse *tko* RNA levels in 2-day old adults of each sex, background, genotype and culture temperature (Fig. 4). At 25 °C the expression level of *tko* was similar in wild-type flies of each sex in the two backgrounds (Fig. 4A), being only slightly higher in Oregon R. We thus used RNA from flies of the corresponding sex and background as a control, in all subsequent assays of *tko* expression in *tko^25t^*flies (Fig. 4B–4D). In all cases tested, *tko* expression was higher in *tko^25t^* than control flies, and the difference was more pronounced in males, where it was greatest in the Oregon R background. These findings suggest that increased *tko* expression might be a compensatory response to (or at least a marker for) a severely deleterious phenotype. However, expression in females was almost identical in the two backgrounds, despite the phenotypic differences and was very similar at the two culture temperatures. Thus, *tko* expression level is not a simple determinant of the *tko^25t^* phenotype.

**Figure 4:**
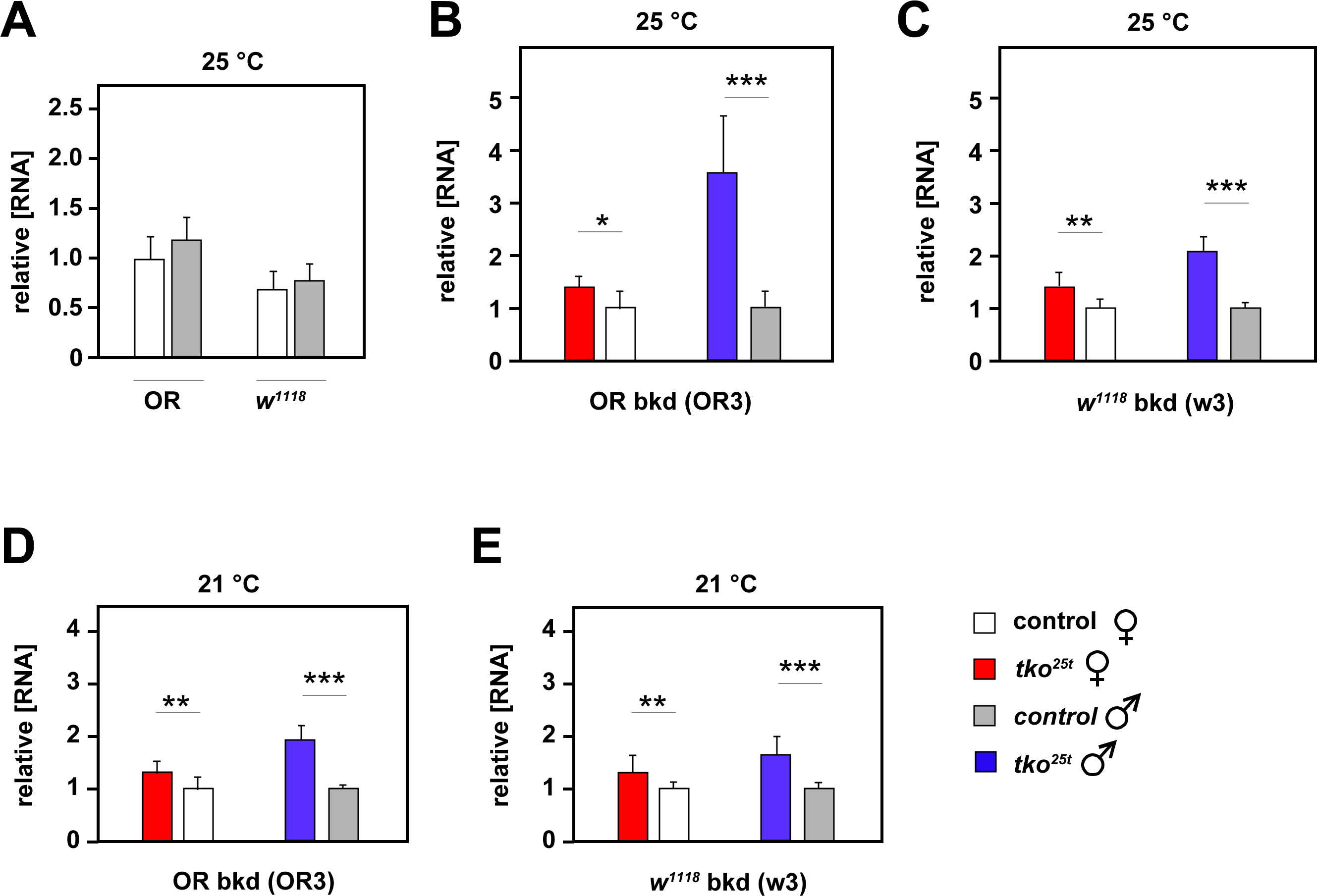
Expression level of *tko* in *tko^25t^* flies backcrossed into two nuclear backgrounds. *tko* RNA levels by qRTPCR (means ± SD), for flies of the indicated sex, background and culture temperature, normalized (A) to values for one isolate of Oregon R wild-type females and (B) to mean values for control flies of the same sex and background (Oregon R wild-type or *w^1118^*, as appropriate). *, **, *** denote significantly different data classes (pairwise comparisons between genotypes, same sex and culture temperature, by Student’s *t* test, p < 0.05, 0.01 or 0.001, respectively). For (A), despite experiment-to-experiment variation, *tko* expression was generally lower in the *w^1118^* background in both sexes and at all temperatures tested, as in the example shown. n = 3 technical replicates of each of 3 biological replicates of batches of 5-7 adults, for all genotypes and sexes.

### Autosomal semi-dominance and parent-of-origin effects underlie strain differences in *tko^25t^* **phenotype**

In an attempt to characterize the genetic basis of the strain background differences we crossed heterozygous *tko^25t^* / FM7 females in each background with *tko^25t^* males in the other, and assessed the phenotype of the first-generation progeny. Since the exact eclosion timing varies slightly between the background strains, and can also differ between experiments, we plotted the amount of developmental delay for both *tko^25t^*females and males in each background, alongside the delay observed for the *tko^25t^*background hybrids (Fig. 5). Female *tko^25t^* hybrid progeny from *tko^25t^* / FM7 mothers in the *w^1118^* background were only slightly more delayed than if both parents were in the *w^1118^* background (Fig. 5A). For males, the delay for hybrid progeny from these mothers was the same as when both parents were in the *w^1118^* background (Fig. 5B). Hybrid *tko^25t^* progeny of both sexes showed an intermediate delay when the female parent was from the Oregon R background (Fig. 5A, 5B). These findings suggest a mixed contribution from autosomal and maternal determinants. The latter could be mitochondrial, although the mtDNA of the two lines was identical at the start of the back-cross. It may also indicate a form of parental imprinting (Lloyd, 2000; Anaka et al. 2009), although the existence of imprinting in *Drosophila* has been questioned (Coolon et al. 2012). Alternatively it could be a classic maternal effect, e.g. involving non-coding RNAs (Soleimani et al. 2020), localized (Hoch and Jäckle 1993) or pioneer transcription factors (Gaskill et al. 2021) or their mRNAs, or other regulatory proteins (Zhang et al. 2018). A simple role for the X-chromosome in mutant eclosion timing seems to be excluded, since the *w^1118^* background appears to be dominant only when inherited from the mother, and in both sexes. Although clear and mostly statistically significant, the differences in delay between the crosses and sexes are too small to enable a meaningful next-generation cross to be conducted, in order to try to tease out the contributions of autosomal, mitochondrial and possible X-chromosomal inheritance.

**Figure 5:**
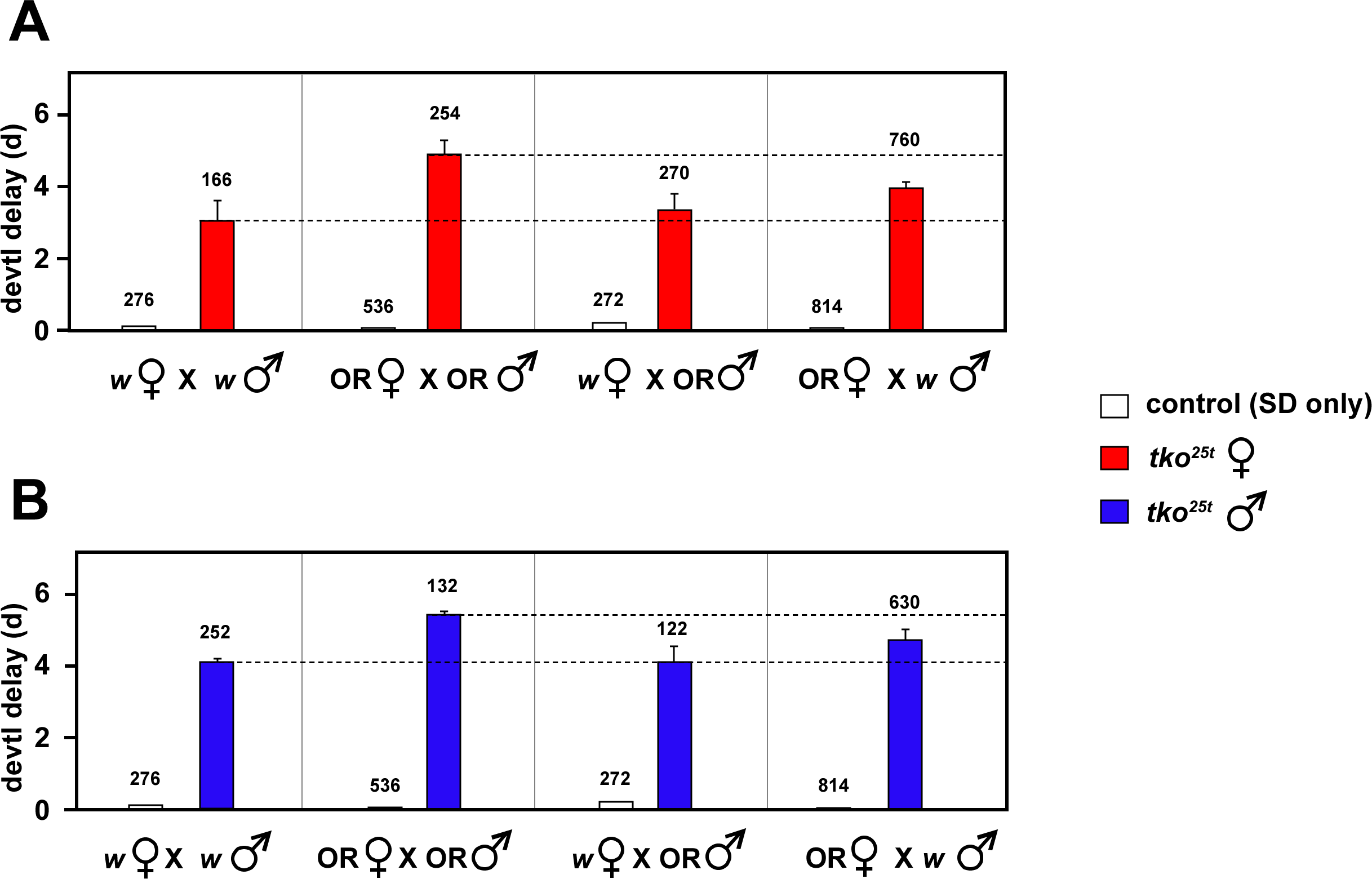
Eclosion timing in *tko^25t^* strain-background crosses. Developmental delay (days, d, means ± SD) at 25 °C of (A) female and (B) male *tko^25t^* progeny from the indicated crosses between *tko^25t^*/ FM7 females and *tko^25t^* / Y males in the backgrounds as shown (OR – Oregon R, *w* – *w^1118^*). For ease of comparison, dotted lines indicate values for the non-hybrid crosses, reproduced from Fig. 3A, which were conducted in parallel using same expanded parental stocks. Data from (A) and (B) are from the same crosses, but with the sexes shown separately, for clarity. Control flies were *tko^25t^* / FM7 heterozygous females, which show an identical eclosion timing as wild-type males (see Fig. S1A), with only the SD shown (means were all set to zero, so as to exclude minor, strain-specific differences in eclosion timing from the analysis). The number of adult flies of each genotype/sex analyzed is indicated above the applicable column. n = 3 replicate bottles in all cases.

We next examined bang-sensitivity in the progeny of these crosses (Fig. 6). Although the recovery times are not normally distributed, making standard statistical tests unreliable, it is clear from the cumulative plots for each sex that a rather similar pattern of dominance and parent-of-origin effects is seen as for developmental delay. In the hybrid crosses, the phenotype of *tko^25t^* progeny from mothers of the *w^1118^* background but Oregon R fathers (green data plots) is very similar to that of *tko^25t^* flies of the pure *w^1118^* background (red data plots). *tko^25t^* females with Oregon R mothers but *w^1118^* fathers (blue data plots, Fig 6A) are intermediate between the two pure lines (red *versus* black data plots), whereas *tko^25t^* males with Oregon R mothers but *w^1118^* fathers (blue data plots, Fig. 6B) are almost identical with *tko^25t^* males of the pure OR background (blue *versus* black data plots, Fig. 6B). Thus, the data indicate an overall ‘dominance’ of the milder *w^1118^* cytoplasm or epigenome, as well as a negative influence on recovery from mechanical shock of the Oregon R X-chromosome.

**Figure 6:**
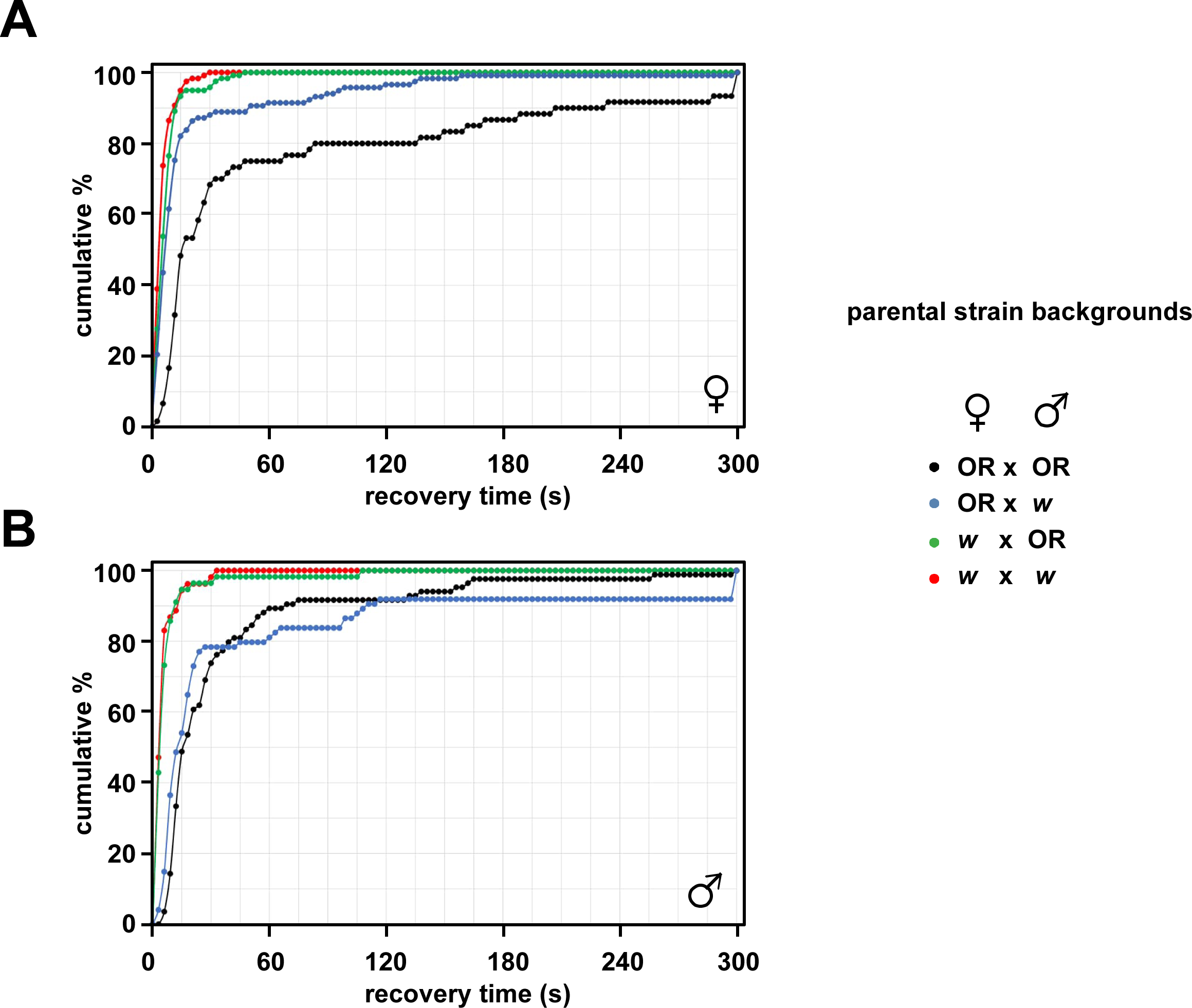
Bang-sensitivity in *tko^25t^* strain-background crosses. Recovery times (plotted cumulatively, over time) at 25 °C for (A) female and (B) male *tko^25t^* progeny from the indicated crosses between *tko^25t^*/ FM7 females and *tko^25t^* / Y males of the indicated parental strain backgrounds (OR – Oregon R background, *w* – *w^1118^* background). Control progeny from all crosses were not bang-sensitive. n = 60 (OR x OR), 117 (OR x w), 119 (w x OR), and 118 (w x w) females, and 84 (OR x OR), 74 (OR x w), 56 (w x OR), and 53 (w x w) males.

### *tko^25t^* courtship analysis is compromised by background effects

We attempted to analyse a third phenotypic parameter of *tko^25t^*flies, namely defective male courtship (Toivonen et al. 2001). However, this did not prove meaningful for the following reasons. Immediately after the back-cross, males in the Oregon R background were extremely weak and unable to mate. Furthermore, previous investigators have found that mutation of the *white* locus itself impairs male courtship behavior, although flies do eventually mate in sufficient numbers to maintain *white^–^*lines. Working on the assumption that the *w^1118^* mutation itself was the cause of the line’s courtship defect, as previous authors have inferred (Xiao et al. 2017),we created a line of Oregon R flies back-crossed into the *w^1118^* background, but retaining red eyes (this was also true of the *tko^25t^* back-cross into *w^1118^*, since *tko* and *w* map very close together on the X-chromosome). Red-eyed *tko^25t^*males in the *w^1118^* background were courtship-impaired (Fig. 7) in a similar manner as inbred *tko^25t^* males studied previously (Toivonen et al. 2001), but red-eyed wild-type males in the *w^1118^* background were also courtship-impaired (Fig. 7), rendering the main experiment meaningless. Although seemingly at odds with the previous findings on the male-courtship effects of *white* (Xiao et al. 2017), it should be noted that the males derived from the back-cross undertaken by the previous authors gave only 6 successful copulations out of 16 trials (Xiao et al. 2017). Furthermore, these authors showed that the absence of eye color *per se* was not the cause of the *w^1118^*courtship defect (Xiao et al. 2017), and that the genotype of females in the assay also influenced the copulation success of *w^1118^* males. In conclusion, although *tko^25t^* males in the Oregon R background were unable to mate, the less severe courtship defect in the *w^1118^* background was impossible to quantify reliably. Note that, despite the male courtship defect of *w^1118^* males, they are not bang-sensitive, as confirmed here (Fig. 2D).

**Figure 7:**
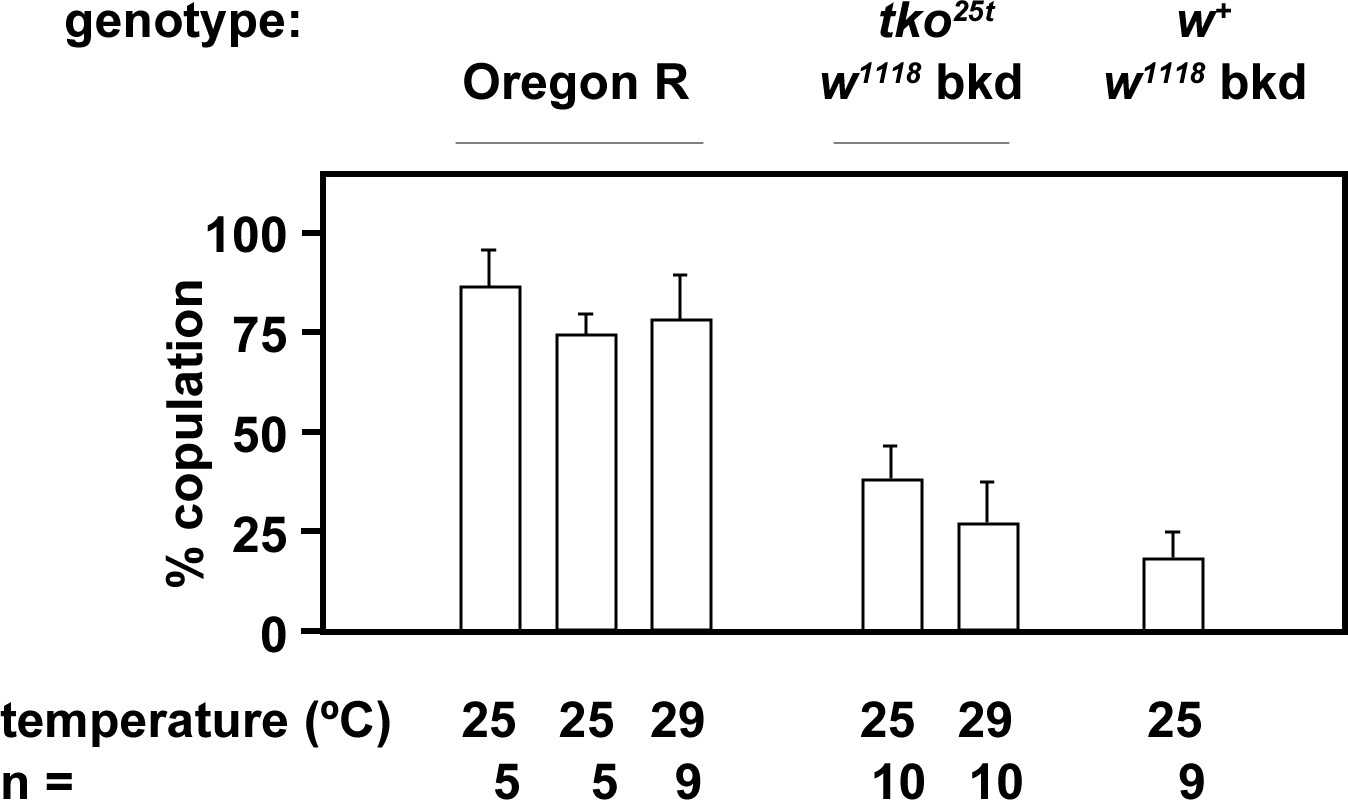
Male courtship effects of *tko^25t^* and *w^1118^*. Percentage of successful copulation (mean ± SEM) of multiple samplings of 6 pairs of flies, at the indicated temperatures. In all assays, male and female flies were of the same genotype and background, and all had red eye color. n – number of batches analyzed for each genotype as indicated. Note that, to exclude the possibility of contamination, the *tko* locus from the red-eyed control flies back-crossed into the *w^1118^* background was sequenced and confirmed to be *tko^+^*.

### Concluding comments

After its initial isolation in 1972, the *tko^25t^* mutant has been maintained in various backgrounds in different laboratories during 50 years, and its exact history is practically impossible to reconstruct. During this time, as a result of inbreeding and inadvertent selection in many different laboratories, the phenotype has become milder (Fig. 2A). A restoration of the original semilethal phenotype was achieved by backcrossing into standard reference strain Oregon R. However, despite being maintained subsequently over a balancer chromosome with no deliberate selection, its phenotype has again become partially alleviated, during 2 years of standard laboratory maintenance. For any mutant strain with a nonlethal phenotype, especially one like *tko^25t^*, where reproductive competence and developmental timing are affected, we would recommend that the strain be periodically backcrossed into the original strain background and the phenotype rechecked, before the strain is used in any experimental context.

Given that the phenotype recorded after back-crossing into two commonly used and broadly wild-type nuclear backgrounds (apart from the *w* mutation in *w^1118^*) was so different (semilethal versus fully viable, 5-10-fold difference in median recovery times from mechanical shock, ∼30% increase in developmental delay, mild male courtship defect versus inability to mate) we would advise to conduct such a back-cross additionally into several other reference strains. Although partial phenotypic suppression of *tko^25t^* by functional absence (or heterozygosity) of *white* cannot be excluded, it seems more likely that other background alleles not knowingly under selection are responsible for the phenotypic differences.

In principle, the novel mutations or background heterogeneity revealed by our study could affect many diverse functions or pathways. The *tko* gene product is part of a complex structure, the mitoribosome, containing dozens of other polypeptides and RNA moieties. Furthermore, the mitoribosome directly influences the functional expression of the 13 mtDNA-encoded polypeptides and, indirectly, the dozens of nuclear-coded polypeptides with which they interact, both transiently and within stable complexes, to maintain essential the biological processes of respiration and oxidative phosphorylation (OXPHOS). In turn, these fundamental metabolic functions affect other key processes, including ionic homeostasis, cell death regulation, steroid synthesis and resistance to oxidative stress, so the instrumental alleles might govern any or many of these. Investigating all of them would be a major task, although some insight might be gained from analyzing the composition, relative abundance and assembly status of the OXPHOS complexes using complexomics (Heide et al, 2012), and from respirometry (Gaviraghi et al, 2021), to try to pinpoint a specific biochemical target in the OXPHOS system. A more classical, genetic approach would be to conduct a mutational screen in the freshly rederived Oregon R background, to look for a suppressor.

This problem of cryptic mutations in inbred reference strains is a widespread phenomenon. The rapid generation time and use of different culture conditions in different laboratories, not to mention possible human error in stock passaging, mean that different isolates of the same reference strain are unlikely to be or to remain genetically identical, and thus may confer different traits upon a given mutation such as *tko^25t^*. The *Drosophila melanogaster* Genetic Reference Panel DmGRP (Mackay et al. 2012), a collection of outbred/inbred fly lines, is an extremely valuable tool in this regard, with a vast array of applications, both potential and realized (e.g., see Spierer et al 2021; Havula et al. 2022). It could be more widely used by the community, to identify background polymorphisms that influence the phenotypic expression of mutations such as *tko^25t^*, especially given the increasing recognition of the importance and flexibility of *Drosophila* in modeling human diseases (Verheyen, 2022). The same issue plagues mouse genetics, for example, where disease-modeling mutations can have no phenotype in one background, yet are lethal in another. The creation of a reference collection of outbred/inbred mouse strains (the CC resource, equivalent to DmGRP) to model complex traits in a genetically diverse population has yielded hundreds of such lines (Churchill et al. 2004; Abu Toamih Atamni et al. 2018).

Ideally, unexpected phenotypes should be verified by back-crossing to a different isolate of the same background strain. Given our previous observations of profound effects of different mtDNA backgrounds on *tko^25t^* (Chen et al. 2012; Salminen et al. 2019), we would recommend that attention should also be paid to mitochondrial genotype, in such back-crossing schemes, regardless of whether the expected phenotype is obviously ‘mitochondrial’. Note that the phenotypic differences observed in our back-crossed *tko^25t^* lines is not the result of genetic heterogeneity within the wild-type lines, even though this may be considerable, as inferred previously for Oregon R (Lints and Bourgois 1987). Back-crossed *tko^25t^* lines within each background had relatively uniform phenotypes.

Given the functional pleiotropy of the *tko* gene product and of the *tko^25t^* mutation, it is hardly surprising that the mutation imposes a selective pressure that can be compensated by diverse changes in genotype or gene expression. But in these respects *tko* is far from unusual. Therefore, the possible influence of genetic background, of selection operating for a supposedly recessive mutation even in the heterozygous condition, and of parent-of-origin effects, should not be discounted for any mutation under study in *Drosophila* or in any model organism.

## DATA AVAILABILITY

All data in this paper is freely available on request. Source data files for each numerical figure are available as spreadsheet tables. Supplementary data files are deposited at Figshare.com as follows: Supplementary Fig. S1 – https://figshare.com/articles/figure/Figure_S1_pdf/22277254, Supplementary Tables S1 and S2 – https://figshare.com/articles/dataset/Supplementary_tables_S1_and_S2/22277287.

## ACKNOWLEDGMENTS

We thank Lotta Kulmala and Eveliina Teeri-Kahelin for technical assistance. The work was conducted with practical support from the Biocenter Finland-funded Tampere Drosophila Facility.

